# Chaperonin facilitates protein folding by avoiding polypeptide collapse

**DOI:** 10.1101/126623

**Authors:** Fumihiro Motojima, Katsuya Fujii, Masasuke Yoshida

**Author notes:** To whom correspondence should be addressed: Fumihiro Motojima, Biotechnology Research Center and Department of Biotechnology, Toyama Prefectural University, 5180 Kurokawa, Imizu, Toyama 939-0398, Japan. present address: Biotechnology Research Center and Department of Biotechnology, Toyama Prefectural University, 5180 Kurokawa, Imizu, Toyama 939-0398, Japan (Fumihiro Motojima). present address: Daiichi Yakuhin Kogyo Co.,Ltd., Kusashima 15-1, Toyama, Toyama 930-2201, Japan (Katsuya Fujii).

## Abstract

Chaperonins assist folding of many cellular proteins, including essential proteins for cell viability. However, it remains unclear how chaperonin-assisted folding is different from spontaneous folding. Chaperonin GroEL/GroES facilitates folding of denatured protein encapsulated in its central cage but the denatured protein often escapes from the cage to the outside during reaction. Here, we show evidence that the in-cage-folding and the escape occur diverging from the same intermediate complex in which polypeptide is tethered loosely to the cage and partly protrudes out of the cage. Furthermore, denatured proteins in the chaperonin cage start their folding from extended conformations but not from compact conformations as usually observed in spontaneous folding. We propose that the formation of tethered intermediate of polypeptide is necessary to prevent polypeptide collapse at the expense of polypeptide escape. The tethering of polypeptide would allow freely mobile portions of tethered polypeptide to fold segmentally. The folding acceleration and deceleration by chaperonin for various substrate proteins can be explained by considering the tethering.

## Introduction

Many cellular proteins require the assistance of molecular chaperones when they fold into the native structure. The system of a bacterial chaperonin, GroEL/GroES, a molecular chaperone that has been studied extensively, assists folding of nascent or denatured proteins in an ATP-dependent manner. GroEL consists of two rings stacked back-to-back. Each ring containing seven 57-kDa subunits forms a large central cavity. GroES is a dome-shaped heptameric ring of 10-kDa subunits. GroEL binds denatured protein at the hydrophobic apical end of the central cavity. Upon binding of ATP to GroEL, GroES attaches to the apical end of GroEL ring as a lid. Then denatured protein is encapsulated into the closed cavity (cage) and starts folding. After several seconds coupled with ATP hydrolysis, the lid GroES is detached and the encapsulated substrate protein, folded or not, becomes free in bulk solution (Hartl & Hayer-Hartl, 2002; Fenton & Horwich, 2003).

Several models have been proposed to explain chaperonin-assisted folding. According to the passive Anfinsen cage model, proteins fold in spontaneous manner in the cage without a risk of aggregate formation (Horwich *et al*, 2009; Ellis, 1994). The iterative annealing model assumes that repetitive binding and release of denatured protein to and from GroEL induces annealing of a misfolded intermediate, thereby enabling its folding into its native state (Stan *et al*, 2003; Shtilerman *et al*, 1999). The confinement model points out that the restriction of the conformational variety of denatured protein in the narrow cage accelerates protein folding by decreasing the activation entropy (Tang *et al*, 2008; Chakraborty *et al*, 2010; Tang *et al*, 2006). Any of the models described above includes the assumption that a whole polypeptide in the cage is enclosed entirely and it is completely isolated from environment. However, our previous studies show that the denatured protein in the cage is accessible by antibody, protease, and trap(D87K) from outside (Motojima & Yoshida, 2010; Motojima *et al*, 2012). Often, it even escapes out of the cage. Actually, polypeptide in the cage is loosely, non-covalently tethered to the interfaces of GroES/GroEL and GroEL subunits, and partially threading out of the cage through the interfaces. Single turnover experiments in which GroES remains bound during the whole folding reaction reveal that a denatured protein in the cage follows two pathways: it folds in the cage (in-cage folding) or it escapes out of the cage, followed by spontaneous folding (out-of-cage folding) (Motojima & Yoshida, 2010; Motojima *et al*, 2012; Motojima, 2015). Herein, we present kinetic evidence for the presence of a common intermediate from which the two pathways diverge. The salient implication is that chaperonin can assist efficient folding by formation of the tethered intermediate, which physically prevents the hydrophobic collapse of the whole polypeptide and allows free segments to undergo productive folding.

## Results

### In-cage and out-of-cage folding diverge from the same intermediate

We have demonstrated that denatured protein in the chaperonin cage folds in the cage (in-cage folding) or escapes out of the cage followed by spontaneous folding in the bulk solution (out-of-cage folding) (Motojima & Yoshida, 2010; Motojima *et al*, 2012). The reaction scheme by which in-cage and out-of-cage folding pathways diverge from the common tethered intermediate has been proposed (Fig. 1). However, other possibilities remain such that each pathway has its own precursor intermediate (Fig. S1). To discern the tethering scheme, we analyzed the kinetics of incage folding and the escape of denatured polypeptide. If this scheme is a typical competitive reaction, its kinetics are described as follows (Equation 1-3).

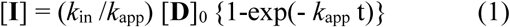

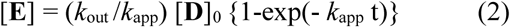

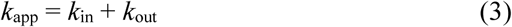

**Figure 1.**
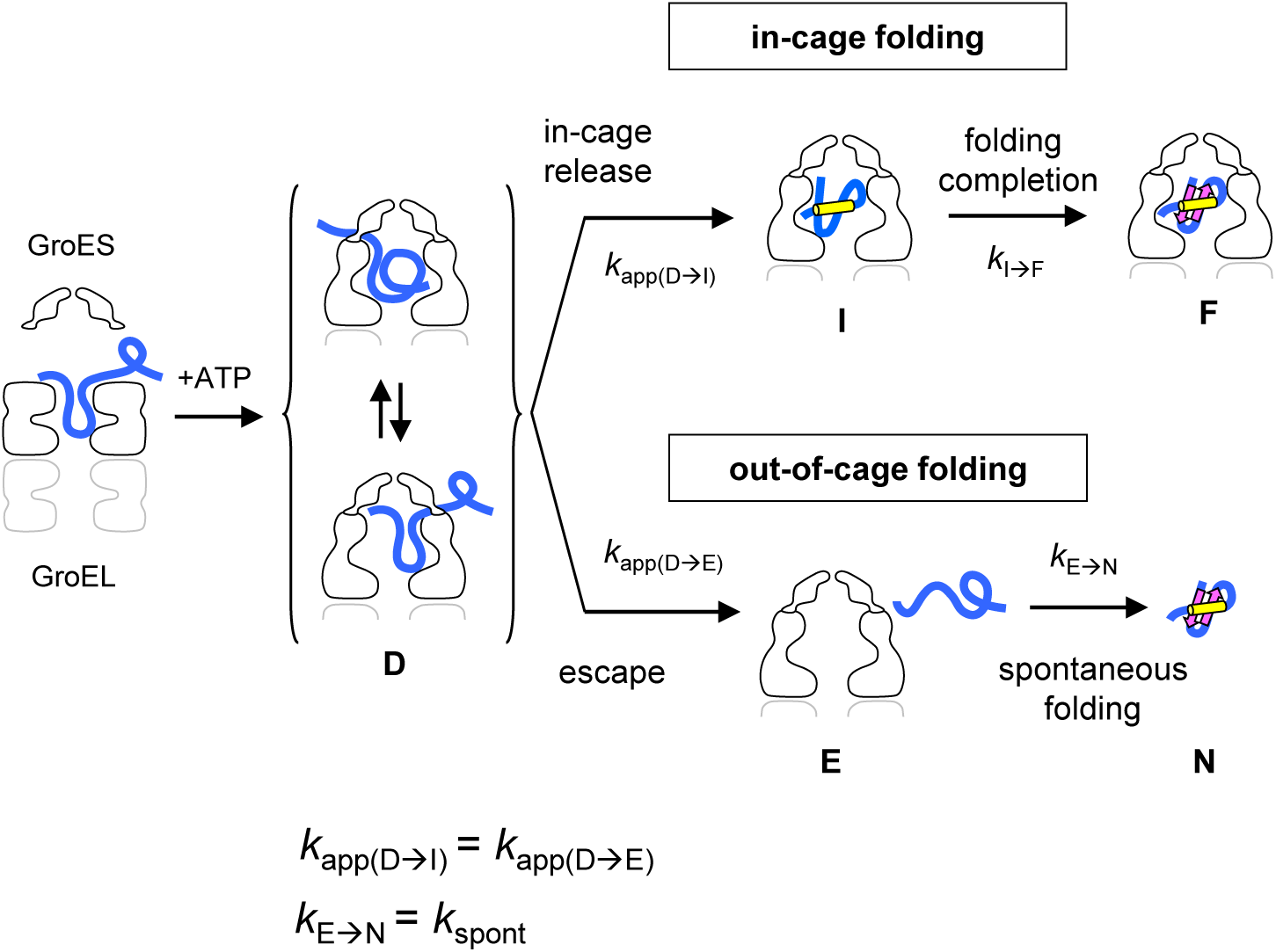
The reaction scheme of chaperonin-assisted protein folding. Tethering scheme for the mechanism of chaperonin-assisted protein folding and kinetic predictions from the scheme. Dissociation of GroES accompanied by ATP hydrolysis is not included in the scheme Tethering intermediate (**D**) is an ensemble of the heterogeneous species in dynamic equilibrium, in which tethering interaction site(s) in the polypeptide chain and in the cage wall are shifting quickly. In-cage folding and out-of-cage folding compete for **D**. In-cage folding includes in-cage release, in which the tethered polypeptide is released into the cage to generate an intermediate (**I**), and completion of folding, in which the polypeptide in the cage gains native structure (**F**). Out-of-cage folding includes escape of the tethered polypeptide out of the cage as a denatured protein (**E**) and folding to native structure in the medium (**N**). If this scheme is really the case, in-cage-release and escape occur apparently at the same rate 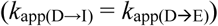, and 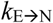 is equal to the rate of spontaneous folding (*k*_spont_).

[**D**]0 stands for the initial concentration of denatured protein bound on GroEL or its single-ring variant. *k*_in_ and *k*_out_ signify an authentic rate constant of the in-cage release and that of the escape, respectively, in the absence of their competing reaction. Their values are obtained from *k*_app_ (= *k*_in_ + *k*_out_) and the fraction of the yield of the in-cage and out-of-cage folding (*k*_in_/*k*_app_ and *k*_out_/*k*_app_, respectively). These equations predict that, as far as the tethering scheme in Fig. 1 is valid, generation of the two products, **I** and **E**, must proceed apparently at the same rate (*k*_app_). If in-cage folding and out-of-cage folding are independent, their apparent rates and yields should be different (Fig. S1). In addition, the rate constant of the transition from **E** to **N** should be equal to that of spontaneous folding (Fig. 1). To test whether these relationships of the rates are really observed, we analyzed the in-cage folding and out-of-cage folding of blue fluorescent protein (BFP). We used a cysteine-less variant of BFP(C49A/C71V), which is designated as BFP hereinafter if not otherwise noted. The spontaneous folding of BFP obeys single exponential kinetics (*k*_spont_ = 0.019 s^−1^), whereas GFP folds in two-phasic manner (Makino *et al*, 1997; Tang *et al*, 2008). To observe the single-turnover folding reaction, we used a single-ring version of GroEL, SR398, which includes a mutation of D398A that makes ATP hydrolysis very slow (Rye *et al*, 1997). Once GroES binds to SR398 upon ATP addition, it remains bound throughout whole period of the folding reaction (Motojima & Yoshida, 2010; Weissman *et al*, 1996; Motojima *et al*, 2012). Regarding gel filtration analysis of the reaction mixture of SR398-assisted folding after folding reactions finished, two BFP fluorescent peaks, the SR398-GroES-BFP ternary complex (45% of total BFP) and free BFP (55%), appeared (Fig. 2a), as observed previously for GFP folding (Weissman *et al*, 1996; Motojima & Yoshida, 2010). The free BFP peak disappeared in the experiment in which trap(D87K) was added immediately before the start of folding reaction, indicating that BFP in a denatured state escapes from the cage and folds spontaneously to produce native BFP in the bulk solution. The time course of the appearance of denatured BFP in the bulk solution was monitored by the increase in FRET efficiency induced by the binding of donor-labeled BFP (BFP(E172C)_Alexa_) to acceptor-labeled trap(D87K) (trap(D87K)_Tx_) in the bulk solution (Fig. 2b, Curve F). After the addition of ATP to start the reaction, FRET efficiency increased with a single exponential function, giving the apparent rate (*k*_app(D→E)_ = 0.042 s^−1^) and leveled off at the escaped fraction 47%. The disappearance of denatured BFP from the cage was monitored using FRET between donor-labeled SR398 (SR398(E315C)_Alexa_) and acceptor-labeled BFP (BFP(E172C)_TMR_) in the absence of trap(D87K) (Fig. 2b, Curve H). Again, the apparent rate *k*_app(D→E)_ = 0.042 s^−1^ was obtained. No initial lag phase exists in the reactions (Fig. S2a), confirming that **D** is an immediate product of encapsulation and that it is a direct precursor of the escape. Thus, disappearance of denatured BFP from the cage and appearance of denatured BFP in the bulk solution take place at the same apparent rate constant of 0.042 s^−1^, which corresponds to *k*_app_ in Equation 2. In these experiments, inclusion of non-labeled trap(D87K) in the mixture did not change the kinetics (Fig. 2b, Curve G), thereby confirming that trap(D87K) did not affect the escape kinetics. Fluorescence labeling used in these experiments did not cause a marked change of folding kinetics of BFP at 25°C (Fig. S2b).

**Figure 2.**
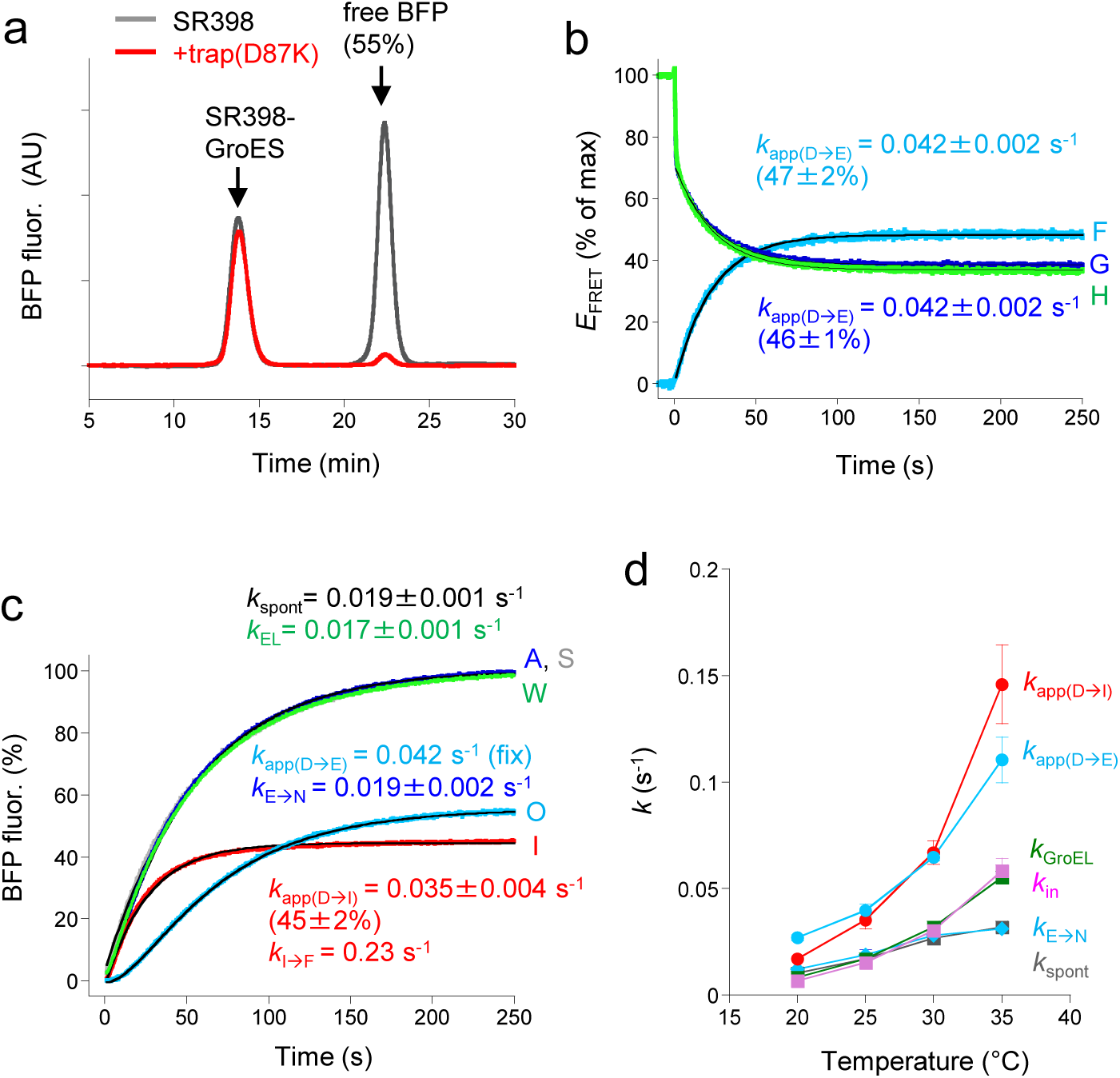
Chaperonin-assisted folding of BFP. (a) Gel filtration analysis of SR398-assisted BFP folding in the absence (SR398, grey) or presence of trap(D87K) (+trap(D87K), red). (b) Time course of the escape of BFP. Curve F, FRET between donor-labeled BFP(E172C)_Alexa_ and acceptor-labeled trap(D87K)_Tx_ shown as percent of maximum FRET efficiency. Curve G and H, FRET between donor-labeled SR398(E315C)_Alexa_ and acceptor-labeled BFP(E172C)_TMR_ in the absence (curve G) or presence of trap(D87K) (curve H) shown as percent of maximum FRET efficiency before ATP addition. The fitting curves giving the indicated rate constants are colored in black. The escape yield of curve G is estimated from the ratio between *E*_FRET_ value at 250 s and that immediately after ATP addition. (c) BFP folding assisted by SR398 at 25°C. Curve A (blue), SR398-assisted folding; curve W (green) GroEL-assisted folding; curve S (grey), spontaneous folding; curve I (red), in-cage folding (folding in the presence of trap(D87K)); and curve O (cyan), out-of-cage folding ([curve A]-[curve I]). The fitting curves giving the indicated rate constants are colored in black. (d) The observed rate constants of BFP folding at various temperatures. *k*_app(D→I)_ and *k*_E→N_ were obtained from BFP fluorescence recovery; *k*_app(D→E)_ was obtained from FRET between BFP(E172C)_Alexa_ and trap(D87K)_Tx_; *k*_GroEL_ is a rate constant of GroEL-assisted folding simulated as a single exponential function; *k*_in_ was calculated from the equation 1. The rate constants averaged from three independent experiments and their standard deviations are shown.

### Direct monitoring of in-cage and out-of-cage folding

BFP folding assisted by SR398 can be monitored directly by the fluorescence recovery of BFP at time resolution ∼0.1 sec, enabling us to determine all the rates of in-cage and out-of-cage folding for the first time. Folding in the presence of trap(D87K) (Fig. 2c, Curve I), which represents in-cage folding, has a short lag (Fig. S2c) and is fitted well by two successive reactions with apparent rate constants: 0.038 s^−1^ and 0.23 s^−1^ (simulations, see Supporting information (SI)). The former value is close to that of the rate of the escape obtained from FRET (0.042 s^−1^). Therefore, it is assumed to be the rate of in-cage release *k*_app(D→I)_. Then, the latter would be the rate of generation of native BFP in the cage (*k*_I→F_). This reaction sequence is opposite from that proposed previously (Ueno *et al*, 2004; Suzuki *et al*, 2008). A value of *k*_I→F_ is more than ten times larger than that of *k*_spont_. BFP released into the cage may not be in a completely unfolded state but has a near-native structure already so that it can gain a native structure very rapidly. The time course of the out-of-cage folding (Fig. 2c, curve O), which was obtained by subtraction of the in-cage folding from the folding in the absence of trap(D87K) ([curve A]–[curve I]), is simulated by two successive reactions with rate constants *k*_app(D→E)_ and *k*_E→N._. When the known value of *k*_app(D→E)_, 0.042 s^−1^, is applied as a fixed parameter, the rate constant *k*_E→N_ = 0.019 s^−1^ gives the best simulation. This value is equal to the rate of spontaneous folding (*k*_spont_ = 0.019 s^−1^). These results all satisfy Equations 1 and 2 and justify the scheme presented in Fig. 1.

### BFP folding at various temperatures

The experiments above mentioned were carried out at 25°C. If the tethering scheme for the chaperonin mechanism is true, relations *k*_app(D→I)_ = *k*_app(D→E)_ and *k*_E→N_ = *k*_spont_ should be observed under any conditions. To examine this, we repeated the same experiments as Fig. 2b and 2c but at different temperatures, 20°C, 30°C and 35°C (Fig. S2d) and the rates were compared (Fig. 2d). The values of *k*_app(D→I)_ and the values of *k*_app(D→E)_ increase in a similar manner as the temperature rises and we consider that they are close enough to meet Equations 1 and 2. Taking a closer look, *k*_app(D→I)_ increases slightly more steeply than *k*_app(D→E)_, probably because of some small bias of the best-fitting simulation of two successive reaction curve I. The values of *k*_E→N_ agree very well with those of *k*_spont_ at all temperatures. The rate constants of BFP folding assisted by wild-type GroEL (*k*_GroEL_) were very close to *k*_in_, but not to *k*_app(D→I)_ of SR398-assisted folding (Fig. 2d), and this reason is also explained by the tethering scheme. In the case of GroEL, out-of-cage folding actually does not occur because an escaped BFP in denatured state is recaptured before it folds spontaneously by other GroEL molecule to enter the next cycle of chaperonin reaction. Without a competing out-of-cage folding reaction, GroEL-assisted folding proceeds only by a manner of in-cage folding at the rates of *k*_in_ and *k*_I→F_. Because the second step of the in-cage folding is very rapid as described, the overall rate of GroEL-assisted folding (*k*_GroEL_) is mostly determined by the first step and is expected to be nearly equal to *k*_in_. This agreement also indicates that the iterative ATPase cycle of GroEL does not accelerate BFP folding. It is interesting that the rate of spontaneous folding is not elevated significantly above 30°C and chaperonin-assisted BFP folding, approximated by *k*_in_, is two times more rapid than spontaneous folding at 35°C. The slow spontaneous folding at 35°C is not attributable to the formation of reversible aggregate because the rate and yield of spontaneous BFP folding at 35°C are unaffected by BFP concentrations (Fig. S2e). Eyring plot of the folding rate constants are shown in Fig. S2f, and the thermodynamic parameters of BFP folding are estimated by fitting with Eyring equation (Supporting Information and Table S1). The fact that chaperonin-assisted folding is slower than spontaneous folding at 20°C indicates that the acceleration of protein folding by chaperonin is dependent on temperature and on the change of activation entropy and enthalpy induced by chaperonin.

### Refolding of protein denatured within the cage

Even if chaperonin-mediated folding proceeds through the tethered intermediate (**D** in Fig.1), a question remains whether **D** is formed only through ATP-induced GroES binding. In the GroEL-denatured protein binary complex, polypeptide of the denatured protein binds to multiple GroEL subunits and it seems plausible that encapsulation is incomplete; one (or more) of the bindings remains as a tether(s) after the binding of GroES to generate the tethered intermediate **D** (Nojima *et al*, 2008). To test this, we observed whether the tethered intermediate **D** is formed when a denatured protein is generated within the cage, but not through GroES binding. As Apetri *et al*. reported for dehydrofolate reductase (Apetri & Horwich, 2008), in-cage denaturation is achieved by exposing a native protein in the cage to a high, but chaperonin-durable, temperature and refolding is initiated by temperature shift-down. We used a single-ring GroEL mutant, SRKKK2, which does not release GroES at 45°C (Fig. S3), and Rubisco as a substrate protein which is denatured at 45°C without escaping from the cage. A native Rubisco monomer folded in the SRKKK2’s cage was heat-denatured completely at 45°C and then refolding was monitored at 25°C. Refolding kinetics of heat-denatured Rubisco in the cage is very similar to that of the control urea-denatured Rubisco with final yields around 100% and rates around 0.12 min^−1^ (Fig. 3). Either for heat-denatured or urea-denatured Rubisco, refolding was inhibited in a short time when trap(N265A) was added to the reaction medium at 1 min after the start of refolding reaction. Trap(N265A) is a strong trap for unfolded polypeptide and it binds to a portion of Rubisco polypeptide protruding out of the cage, thereby preventing further refolding (Motojima & Yoshida, 2010). These results show that the generation of the tethered intermediate is not a result of incomplete encapsulation but an inherent reaction of chaperonin when unfolded polypeptide is present in the cage.

**Figure 3.**
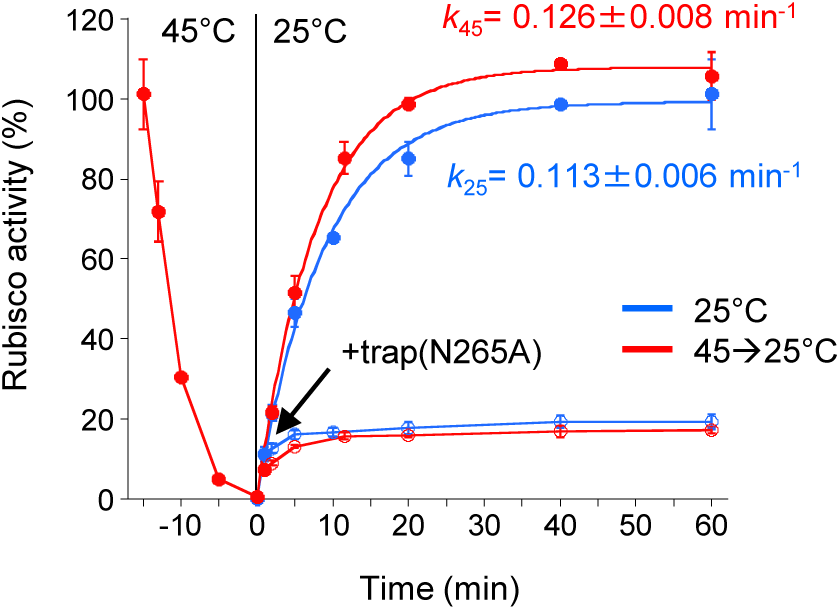
Refolding of Rubisco denatured within the cage. Blue circles: Folding of urea-denatured Rubisco monomer was started by ATP-induced GroES binding to SRKKK2 at 25°C. Red circles: A native Rubisco monomer in the cage of SRKKK2 was heat-denatured by incubation at 45°C (-15 to 0 min), and refolding in the cage was started by a temperature shift to 25°C. Open circles: The folding of Rubisco after addition of trap(N265A) (2 μΜ) at 1 min. Standard deviations from three independent experiments are shown. Other experimental details are described in Material and Methods.

### Tethering prevents polypeptide collapse in BFP folding

The conformational state of BFP during folding was probed using single-pair FRET. Cysteine residues of BFP(K52C/E172C) (Fig. 4a) were labeled by donor (Alexa) and acceptor (tetramethylrhodamine) fluorescent dyes and the change of FRET efficiency was monitored (see details in Methods). Depending on the distance of the two dyes, FRET efficiency becomes minimum when BFP is denatured in acid (*E*_FRET_ = 0.32 ± 0.01) and it becomes maximum when BFP is in a native state in bulk solution (*E*_FRET_ = 0.96 ± 0.01) and in the cage (*E*_FRET_ = 0.95 ± 0.01) (Fig. S4a). In spontaneous folding, the initial FRET efficiency was 0.86 ± 0.02, which increased slowly to 0.95 of the native state at the rate 0.015 s^−1^ (Fig. 4b). This high initial *E*_FRET_ value suggests that, as observed in spontaneous folding of many proteins (Agashe *et al*, 1995; Daggett & Fersht, 2003; Dobson & Karplus, 1999; Walters *et al*, 2013), the freely extending polypeptide of denatured BFP collapses to compact conformations immediately at the intial of folding reaction. In contrast, and in support of the tethering scheme, the initial FRET efficiency of the SR398-assisted folding reaction (*E*_FRET_ = 0.73 ± 0.02) is lower than that of spontaneous folding, indicating that tethered polypeptide is extended. As the tethered BFP is released into or outside of the cage and folds to the native structure, the FRET efficiency increases to the level of native BFP. Similar results were obtained from another FRET pair K3C/239C (Fig. 4c).

**Figure 4.**
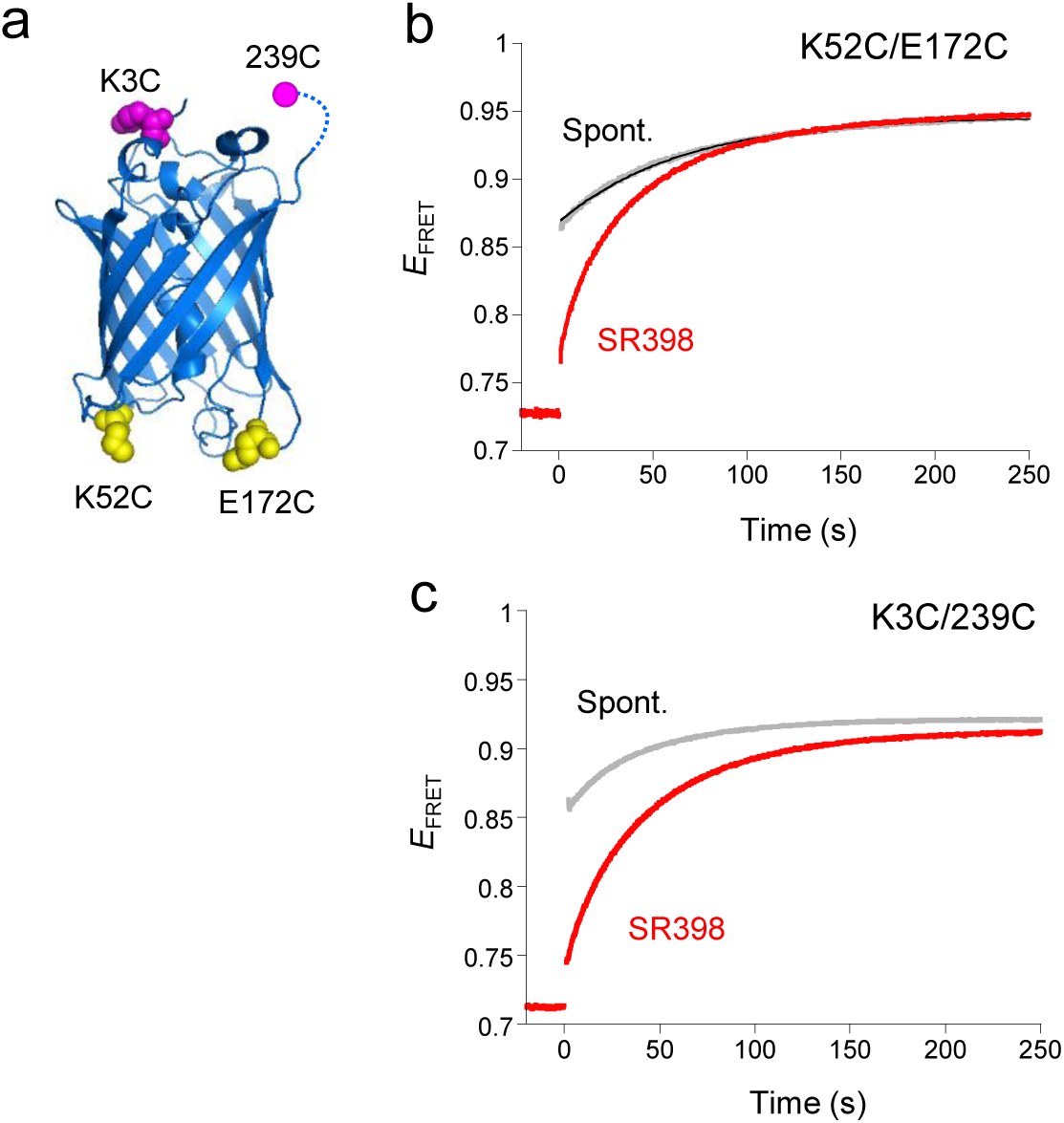
Conformational change of BFP in chaperonin-assisted folding. (a) The residues mutated to cysteine for single-pair FRET. Crystal structure (PDB ID: 1GFL) of GFP is shown in cartoon model. The Cα atoms of cysteine mutations of K52C/E172C and K3C/239C are shown in yellow and magenta CPK models, respectively. (b, c) BFP folding monitored by the change of single-pair FRET efficiency (*E*_FRET_) between fluorescent dyes labeled at K52C/E172C (b) and K3C/239C (c). Spontaneous folding and SR398-assisted folding are shown in grey and red curves, respectively. Reactions were started at time zero by dilution of denatured BFP in spontaneous folding or by ATP-addition in SR398-assisted folding.

### Folding of DMMBP obeys the tethering scheme

We examined next the folding of a double-mutant (V8G/Y283D) of maltose binding protein (DMMBP) (Fig. 5a). DMMBP has been known as a typical protein of which folding is accelerated by chaperonin several times compared to spontaneous folding (Tang *et al*, 2006; Chakraborty *et al*, 2010; Motojima *et al*, 2012). Time course of the escape of DMMBP from the chaperonin cage was monitored by the decease of FRET efficiency between SR398(E315C)_Alexa_ and DMMBP(A52C)_Cy3_, and *k*_app(D→E)_ = 1.04 min^−1^ was obtained (Fig. 5b). After immediate FRET decrease upon ATP addition, about 60% of DMMBP(A52C)_Cy3_ escaped out of the cage. Inclusion of trap(D87K) in the solution did not change the escape kinetics. In-cage-folding of DMMBP(A52C)_Cy3_ was monitored by the recovery of tryptophan fluorescence of native DMMBP in the presence of trap(D87K) (Fig. 5c). Consistent with the escape fraction, the in-cage-folding yield was 42%. The curve of fluorescence increase is well simulated by a single exponential function and an apparent rate 1.04 min^−1^ was estimated. It is likely that the step **I**→**F** is too rapid to be observed and the value of 1.00 min^−1^ represents *k*_app_(D→I). Resemblance of the values of *k*_app(D→I)_ and *k*_app(D→E)_ of DMMBP(A52C)_Cy3_ indicates that SR398-assisted DMMBP(A52C)_Cy3_ folding obeys the tethering scheme. The value of *k*_in_ (0.42 min^−1^) estimated from *k*_app(D→I)_ is higher than *k*_GroEL_ (0.26 min^−1^), suggesting that released DMMBP(A52C)_Cy3_ is not rapidly recaptured and/or refolded by chaperonin. To probe the conformational state during folding, a cysteine pair of DMMBP(D184C/K362C), which are located closely (Chakraborty *et al*, 2010), was labeled with donor and acceptor fluorescent dyes and single-pair FRET was monitored (Fig. 5d and Fig. S4b). In this experiment, we used SR1, a single ring version of GroEL with slow ATP hydrolysis, which we confirmed the same kinetic behaviors as SR398 in these experiments. The FRET efficiency of GroEL-assisted folding of labeled DMMBP increases at a rate (0.24 min^−1^) that is similar to *k*_GroEL_ obtained from tryptophan fluorescence (0.26 min^−1^). In contrast to a previous report (Chakraborty *et al*, 2010), and consistent with the tethering scheme, DMMBP in the cage of GroEL starts folding from a more extended conformation than in the case of spontaneous folding, as indicated by a smaller value of FRET efficiency (0.57±0.01) than the value of the spontaneous one (0.61±0.01) (Fig. 5d).

**Figure 5.**
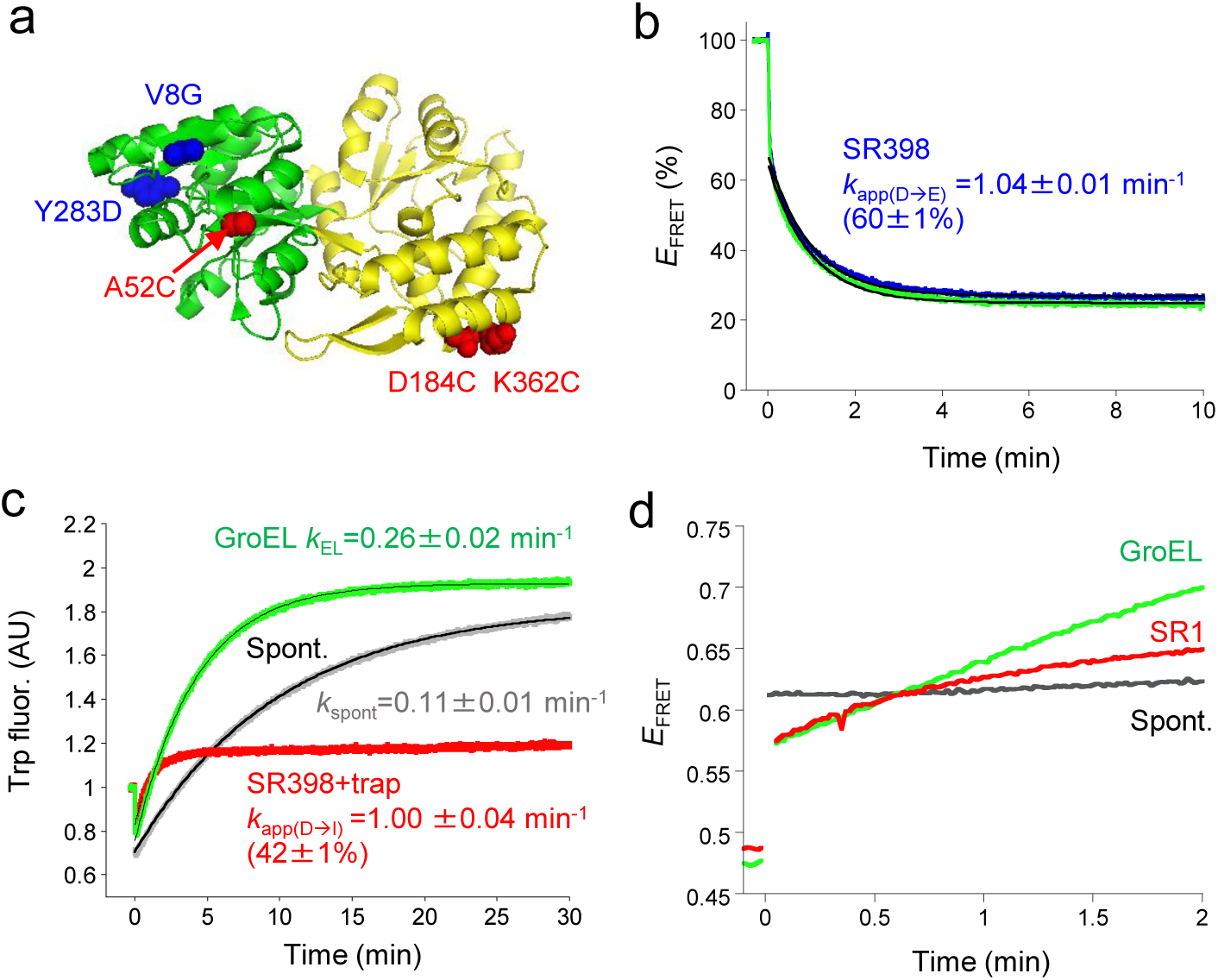
Chaperonin-assisted folding of DMMBP. (a) Mutated residues of DMMBP used for this study. The crystal structure of MBP (PDB ID: 1OMP) is shown in cartoon model. The double-mutation (V8G/Y283D) and cysteine mutation (A52C, D184C, K362C) are colored in blue and red, respectively. (b) Time course of the escape of DMMBP. FRET between donor-labeled SR398(E315C)_Alexa_ and acceptor-labeled DMMBP(A52C)_Cy3_ in the absence (blue) or presence of trap(D87K) (green) are shown as percent of maximum FRET efficiency before ATP addition. The fitting curves giving the indicated rate constants are colored in black. The yield of in-cage folding shown in parentheses was estimated from *E*_FRET_ at time zero (70%). (c) DMMBP folding monitored by recovery of tryptophan fluorescence of native DMMBP. Folding of urea-denatured DMMBP(A52C)_Cy3_ was observed. Spontaneous folding, GroEL-assisted folding, and in-cage folding assisted by SR1 in the presence of trap(D87K) are shown in grey, green, and red, respectively. The in-cage folding yield is estimated based on the observation that magnitude of tryptophan fluorescence of native DMMBP in the chaperonin cage is 78% of that of free native DMMBP(Motojima *et al*, 2012). (d) DMMBP folding monitored by the change of single-pair FRET efficiency (*E*_FRET_) between fluorescent dyes labeled at D184C/K362C. Spontaneous folding, SR1-assisted folding, and GroEL-assisted folding are shown in grey, red, and green, respectively. Reactions were started at time zero by dilution of denatured BFP in spontaneous folding or by ATP-addition in SR398-assisted folding.

### Folding of DapA also obeys the tethering mechanism

DapA is a typical obligate substrate protein in *Escherichia coli* that requires GroEL and GroES for its folding (Georgescauld *et al*, 2014; Kerner *et al*, 2005; Fujiwara *et al*, 2010). DapA mutant containing C20A and a cysteine pair of T3C and E223C, which are located close in the native structure (Fig. 6a), were labeled with the donor (Alexa350) and acceptor fluorescence dyes (Atto488) for single-pair FRET. Escape kinetics observed by the increase of FRET of labeled Atto488 dye on DapA(C20A/T3C/E223C) induced by its binding to trap(D87K)_Tx_ show that 18% of DapA escape out of the cage at an apparent rate *k*_app(D→E)_ of 5.7 min^−1^ (Fig. 6b). Under the condition of spontaneous folding, dilution of urea-denatured DapA immediately produces a rather compact conformational state with single-pair FRET efficiency of 0.66±0.01 and further change does not occur (Fig. 6c). In SR1-assisted DapA folding, folding of labeled DapA started from a state with FRET efficiency of 0.60±0.01, indicating more extended state than spontaneous condition. As folding proceeds, the increasing FRET efficiency approaches a value of native DapA (0.82±0.01) (Fig. S4c). Because labeled DapA is unable to fold spontaneously (Georgescauld *et al*, 2014), the FRET increase mostly reflects in-cage folding and its apparent rate, 4.8 min^−1^, corresponds to *k*_app(D→I)_. Therefore, the values of *k*_app(D→I)_ and *k*_app(D→E)_ are close enough, though not the same, to justify the tethering scheme. The state formed under spontaneous folding condition is competent as a substrate protein for chaperonin-assisted folding; by addition of GroEL, FRET efficiency shifts down to 0.56±0.01 and subsequent addition of GroES/ATP induces an increase of FRET efficiency obeying almost identical kinetics observed for the control GroEL-assisted folding (Fig. S4c).

**Figure 6.**
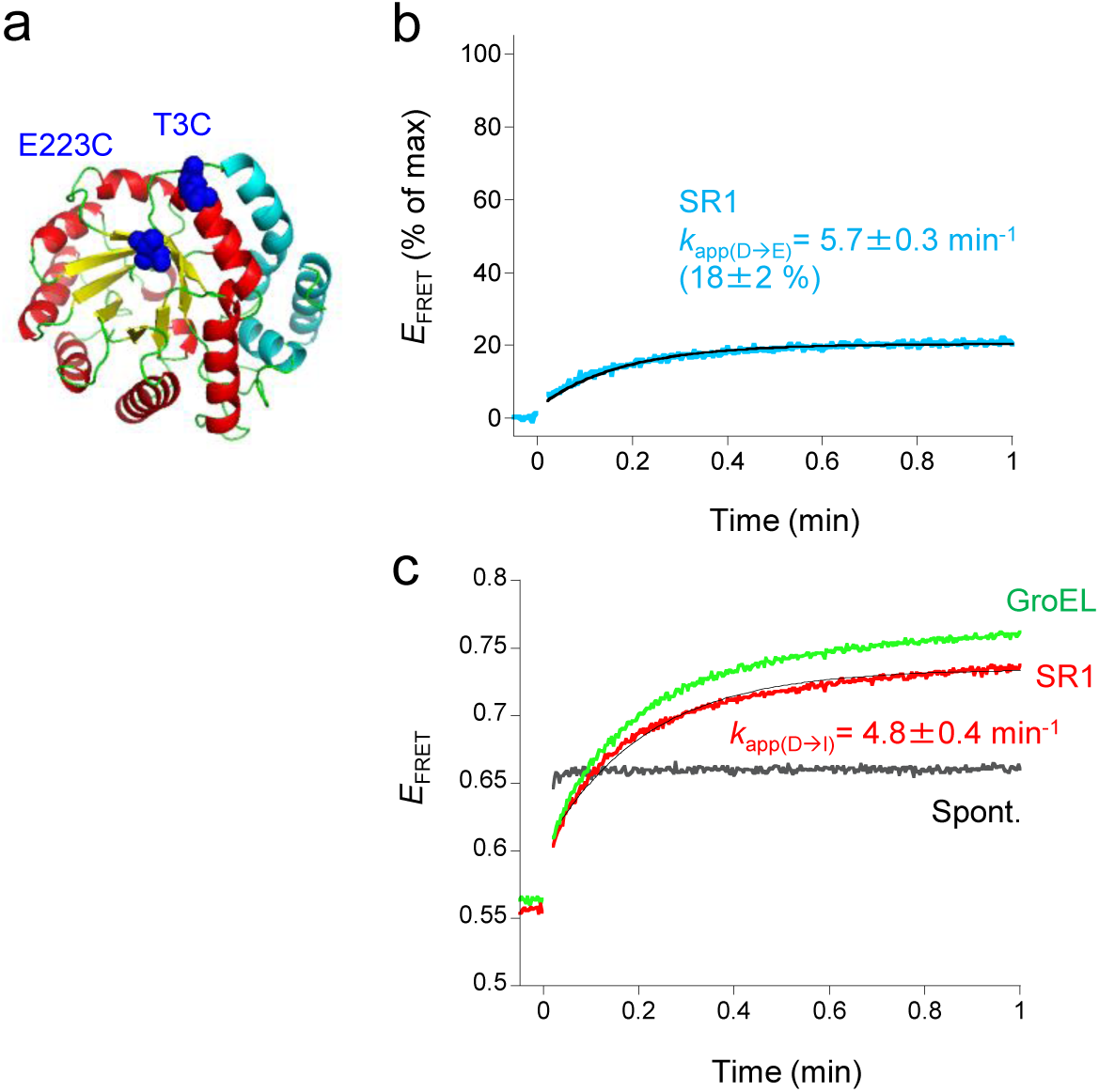
Chaperonin-assisted folding of DapA. (a) Mutated residues of DapA used for single-pair FRET. The monomer structure of DapA is shown in cartoon model (PDBID: 1DHP). T3 and E223 mutated to cysteine are shown in blue CPK model (b) Time course of the escape of DapA. FRET between donor-acceptor labeled DapA(C20A/T3C/E223C) and trap(D87K)_Tx_ are shown as percent of maximum FRET efficiency. The fitting curve giving the indicated rate constant is colored in black. (c) DapA folding monitored by change of single-pair FRET efficiency (*E*_FRET_) between fluorescent dyes labeled at T3C/E223C. Spontaneous folding, SR1-assited folding, and GroEL-assisted folding are shown in gray, red, and green, respectively. The fitting curve giving the indicated rate constant is colored in black.

## Discussion

We have proposed that the in-cage folding and the escape of denatured polypeptide occur from the tethered intermediate in the chaperonin-assisted protein folding (Fig. 1). However, an alternative model is possible that two types of the initial tethering intermediates are formed upon encapsulation of the substrate protein each leading independently to the in-cage folding and the escape (Fig. S1). According to this model, the escape is a result of faulty encapsulation. The tethering mechanism in Fig. 1 contains two reactions competing for the common precursor and kinetic equations of such reaction scheme indicate that the apparent rates of the two competing reactions must be the same. On the contrary, the model shown in Fig. S1 does not require any relation between rate of the in-cage release and rate of the escape. Our observations show that apparent rate constant of the in-cage release and that of the out-of-cage release (escape) of BFP in chaperonin-assisted folding reaction are always sufficiently close to meet the equations. This relation of the apparent rate constants is observed also for DMMBP and DapA. Previous analysis of chaperonin-assisted folding of rhodanese also indicates similar relationships between the two apparent rate constants (*k*_app_ = 0.11 min^−1^, *k*_app(D→E)_ = 0.10 min^−1^) (Motojima & Yoshida, 2010). The kinetic analysis, therefore, provides evidence for the tethering reaction scheme shown in Fig. 1. Furthermore, as shown by generation of the tethered intermediate **D** from native Rubisco monomer in the cage by heat treatment (Fig. 3), tethering occurs by itself whenever a denatured protein exists in the cage. The thermodynamics of BFP folding is also consistent with the tethering of denatured polypeptide by the cage. The increase of the activation enthalpy of in-cage folding of BFP compared with spontaneous folding can be attributed to the interactions between denatured BFP and the cage wall. The increase of the activation entropy can be explained by restricted conformations of denatured protein in the cage.

Another finding in this report is that the conformational state of denatured proteins in the tethered intermediate is more extended than that of spontaneous folding. Actually, within the time resolution limit of the single-pair FRET experiment, folding in the chaperonin cage starts from an extended conformation, whereas spontaneous folding starts from a compact conformation (Fig. 7). This tendency was also observed in Rubisco folding, although compaction of Rubisco upon GroES binding has been focused (Lin & Rye, 2004; Weaver & Rye, 2014). The extended conformation is consistent with the tethering model, but not with previous assumption that the encapsulated protein is free or repulsive to the cage. The compact conformation observed in the spontaneous folding reflects the collapsed state, and thus, protein folds in the chaperonin cage by avoiding initial collapse of polypeptide chain. It is thought that the collapsed state tends to fall into kinetically-trapped state containing nonnative interactions at local energy minimum (Agashe *et al*, 1995; Daggett & Fersht, 2003; Dobson & Karplus, 1999; Walters *et al*, 2013). It should be noted, however, that chaperonin-assisted folding of rhodanese and mouse dihydrofolate reductase are slower than their spontaneous folding (Hofmann *et al*, 2010; Motojima & Yoshida, 2010; Sirur & Best, 2013). In such cases, tethering manner may not be suitable (too strong, too many tethered positions, or unfavorable for productive partial folding, etc.) and productive folding would be slowed. Indeed, the increase of hydrophobic surface in the chaperonin cage by the labeling of hydrophobic fluorescence dyes decreases the folding rate of rhodanese, but inhibits the escape (Motojima & Yoshida, 2010). At the expense of possible escape of polypeptide out of the chaperonin cage, loose tethering is necessary to extend polypeptide and not to decelerate folding too much. When native interactions are stronger than the tethering, the folding rate is not changed by chaperonin as observed in the folding of wild-type MBP (Tang *et al*, 2006). This folding manner is recognized as passive Anfinsen cage model.

**Figure 7.**
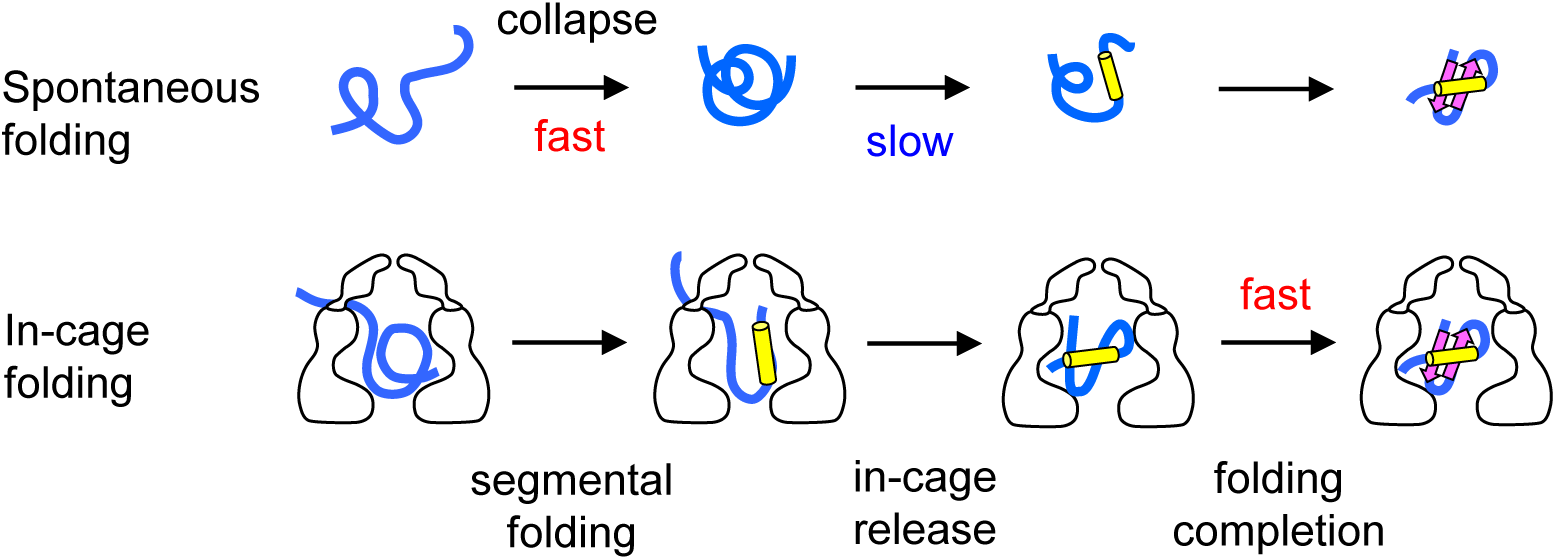
Difference between spontaneous and in-cage folding. In spontaneous folding, denatured protein collapses rapidly to a compact form and tuning of intramolecular interactions to native conformation follows. In-cage folding, on the contrary, denatured protein is dynamically tethered to the chaperonin cage wall with more or less extended form and thus avoids the initial collapse. Segmental folding of free portions of polypeptide away from the tethered position follows and near-native conformation is already formed until the time of in-cage release.

By virtue of real-time monitoring of BFP folding, we clarify that the in-cage folding proceeds through two steps; in-cage release of the tethered protein and folding completion of the released protein to native structure, in contrast to the single step in spontaneous folding. This indicates that the folding pathway of the in-cage folding is different from that of spontaneous folding, even though the overall folding rate of the in-cage folding of BFP is same as that of the spontaneous folding at 25°C. The difference in these thermodynamic parameters also support this notion. It is important to note that the second step is much more rapid than spontaneous folding. It appears that the tethered substrate protein gains a near-native structure by the time it is untethered, leaving only the final step of folding as the second step. In noncovalent tethering, tethered position is shifting quickly and there may be a moment when polypeptide is tethered to a position suitable for the freely mobile portion of polypeptide to generate native-like partial structure. The partial structures thus formed may have some stability and tend to accumulate. Then, free polypeptide region to be tethered becomes narrowed and finally whole protein becomes free to finish folding. In support of this, we observed previously similar rapid folding completion in the cage upon untethering by reduction of rhodanese covalently-tethered to the cage wall through disulfide (Motojima & Yoshida, 2015). As a rapid folding fraction increases depending on the incubation time before untethering, it is clear that partial, productive folding proceeds with time even in the tethered polypeptide. A recent extensive study reveals efficient, sequential progress of segmental folding of DapA in the chaperonin cage (Georgescauld *et al*, 2014). Certainly, this observation reflects the segmental folding of the tethered polypeptide in the cage.

## Supporting information

Supporting information include six figures, two tables, Supporting materials and methods, and Supporting references can be found with this article online.

## Acknowledgements

We are grateful to Y. Ishizaki and Y. Motojima-Miyazaki for protein preparation. This work was supported by MEXT-supported Program for the Strategic Research Foundation at Private Universities (2011-2016), and Grant-in-Aid for Scientific Research (C) (No. JP15K06983 to F. M.).

## Materials and Methods

### Proteins

The mutants of proteins used in this study were prepared using QuikChange multi Lightning Site-Directed Mutagenesis Kit (Agilent Technologies Inc.) with appropriate oligonucleotides. The mutants of GroEL, GroES, and fluorescently labeled GroES were prepared as described (Motojima & Yoshida, 2010; Motojima *et al*, 2012; Motojima & Yoshida, 2003). Cysteine-less BFP was prepared by the mutations of C49A, F65L, Y67H, and C71V in wild-type GFP. MBP, DMMBP, and DapA were prepared as described (Motojima & Yoshida, 2010; Motojima *et al*, 2012; Georgescauld *et al*, 2014).

### BFP folding assays

The reaction mixture containing 0.1 μM GroEL or SR398 and 0.5 μM GroES in buffer HKM (50 mM HEPES-NaOH, pH 7.5, 50 mM KCl, 10 mM MgCl_2_, and 1 mM DTT) was incubated with stirring in a quartz cuvette placed in a water jacket with circulating water at the indicated temperature. BFP was denatured by adding an equal volume of 200 mM glycine-HCl pH 2.5. Denatured BFP (10 μM) was added to the reaction mixture at the final concentration of 0.05 μM. For fluorescently labeled BFP, the final concentration was 0.02 μM. ATP was added to start chaperonin-assisted folding. To measure BFP folding in the chaperonin cage, 0.1 μM trap(D87K) was mixed before ATP addition to eliminate free denatured BFP that had escaped from the cage to the bulk solution. BFP folding was monitored by the fluorescence emission of 440 nm excited at 380 nm using a fluorometer (FP-6500; Jasco Corp.). In rapid-mixing experiment (SFM-400; Biologic), 1.0 μM GroES was used.

### Rubisco folding assay

Rubisco was denatured in urea as reported (Motojima *et al*, 2012). Denatured Rubisco (final 1 μM) was diluted by 50-fold into buffer H5KM (50 mM HEPES-NaOH, pH 7.5, 5 mM KCl 10 mM MgCl_2_, and 1 mM DTT) containing 2 μM SRKKK2, 4 μM GroES, 10 mM glucose, and 10 mM NaHCO_2_. The folding reaction was started by addition of ATP (final = 1 mM). At 10 s after ATP addition, excess ATP was hydrolyzed to ADP by addition of hexokinase (final = 0.01 U/μl). For the experiment of refolding of heat-denatured Rubisco within the cage, the SRKKK2 containing native Rubisco monomer in the cage produced as described above was incubated at 45°C for 15 min. For rapid temperature shift from 45°C to 25°C, the reaction solution was mixed into a micro tube containing the equal volume of buffer H5KM preincubated at 25°C. Generation of native Rubisco monomer was assayed as follows. Aliquots (20 μl) were mixed with ice-cold quenching buffer (40 μl) containing 20 mM Tris-HCl pH7.5, 15 mM EDTA, 1 mM DTT, and 0.5 μΜ trap(D87K). BSA was subsequently added to aliquots (final 0.2 mg/ml) and frozen by liquid nitrogen. The thawed samples on ice were activated and their Rubisco activity were measured by coupled enzyme assay as reported (Motojima *et al*, 2012).

### Gel filtration analysis of BFP

The folding reaction mixture was applied to Superdex 200 10/300 equilibrated with buffer containing 20 mM Tris-HCl pH 7.5, 5 mM KCl, 10 mM MgCl_2_, and 1 mM DTT. Fluorescence of BFP was monitored using an in-line fluorometer (FP-2020; Jasco Corp.).

### FRET observation to monitor the escape

The FRET efficiency change of a donor-labeled substrate protein upon binding to trap(D87K) labeled with TexasRed-maleimide (trap(D87K)_Tx_) was measured as reported (Motojima & Yoshida, 2010). Substrate proteins were denatured as described (Motojima *et al*, 2012; Georgescauld *et al*, 2014). Denatured substrate protein (final = 0.02 μM) was added to the buffer HKM containing 0.05 μM GroEL or 0.1 μM SR-variant at 25°C. When DapA was a substrate protein, 10 mM pyruvate was added and incubated at 20°C. Then, 0.5 μM GroES, 1 mM ATP, and 0.1 μM trap(D87K)_Tx_ for donor fluorescence change in the presence of acceptor TexasRed dye (*F*_DA_) or non-labeled trap(D87K) for donor fluorescence change in the absence of acceptor (*F*_D_) were added to start the assisted folding reaction. The percentage of FRET change was calculated from division of FRET efficiency (1-*F*_DA_/*F*_D_) by maximum FRET efficiency that was obtained when donor-labeled substrate proteins directly bound to trap(D87K)_Tx_. In the case of BFP and DMMBP, the FRET efficiency change of donor-labeled SR398(E315C) (SR398(E315C)_Alexa_, 0.05 μM) and acceptor-labeled substrate protein (0.1 μM) upon release out of the cage of SR398(E315C)_Alexa_ was also measured. In rapid-mixing experiments, 1.0 μM GroES was used.

### Fluorescence labeling for single-pair FRET

For single-pair FRET, we prepared mutants containing only two reactive cysteine residues that were located near each other in the native structure. Labeled proteins, donor dyes and acceptor dyes were the following: BFP(K52C/E172C) and BFP(K3C/239C), Alexa 488-maleimide (Invitrogen) and tetramethylrhodamine-maleimide (Invitrogen); DMMBP(D184C/K362C) and DapA(C20A/T3C/E223C), Alexa350-maleimide (Invitrogen) and ATTO488-maleimide (ATTO Tec.). First, donor fluorescence dye containing maleimide moiety was mixed into protein at 1:2 molar ratio and incubated for 1 h at 25°C. The donor-labeled protein was purified using gel filtration (Sephadex G-25, GE healthcare) to remove free fluorescence dye. Half of the purified product was reserved as donor-labeled sample. The other half of the donor-labeled protein was mixed with four-fold molar excess of the acceptor fluorescence dye containing maleimide moiety. After incubation for 16 h at 4°C, free fluorescence dye was removed using gel filtration to purify donor–acceptor-labeled protein. Single-pair FRET was measured at 25°C (BFP and DMMBP) or at 20°C (DapA). In the case of DapA, the reaction mixture contained 10 mM pyruvate. Denatured proteins were diluted into the buffer and refolded as described above. In case of DapA, to inhibit FRET among DapA subunits in native tetramer, the labeled DapA was mixed with the nine-fold molar of non-labeled DapA and denatured. FRET efficiency (*E*_FRET_) was calculated from the two fluorescence measurements using donor-labeled protein (*F*_D_) and donor–acceptor-labeled protein (*F*_AD_) using the function (*E*_FRET_ = 1 - (*F*_DA_/*F*_D_)).

